# Can Carbon Nanofibers Affect Anurofauna? Study Involving Neotropical *Physalaemus cuvieri* (Fitzinger, 1826) Tadpoles

**DOI:** 10.1101/2021.02.16.431548

**Authors:** Abraão Tiago Batista Guimarães, Fernanda Neves Estrela, Aline Sueli de Lima Rodrigues, Rafael Henrique Nóbrega, Ives Charlie-Silva, Guilherme Malafaia

**Author notes:** Corresponding Author: Biological Research Laboratory, Goiano Federal Institution – Urutaí Campus. Rodovia Geraldo Silva Nascimento, 2,5 km, Zona Rural, Urutaí, GO, Brazil. CEP: 75790-000.

## Abstract

Although carbon nanotubes’ (CNTs) toxicity in different experimental systems (*in vivo* and *in vitro*) is known, little is known about the toxic effects of carbon nanofibers (CNFs) on aquatic vertebrates. We herein investigated the potential impact of CNFs (1 and 10 mg/L) by using *Physalaemus cuvieri* tadpoles as experimental model. CNFs were able to induce nutritional deficit in animals after 48-h exposure to them, and this finding was inferred by reductions observed in body concentrations of total soluble carbohydrates, total proteins, and triglycerides. The increased production of hydrogen peroxide, reactive oxygen species and thiobarbituric acid reactive substances in tadpoles exposed to CNFs has suggested REDOX homeostasis change into oxidative stress. This process was correlated to the largest number of apoptotic and necrotic cells in the blood of these animals. On the other hand, the increased superoxide dismutase and catalase activity has suggested that the antioxidant system of animals exposed to CNFs was not enough to maintain REDOX balance. In addition, CNFs induced increase in acetylcholinesterase and butyrylcholinesterase activity, as well as changes in the number of neuromats evaluated on body surface (which is indicative of the neurotoxic effect of nanomaterials on the assessed model system). To the best of our knowledge, this is the first report on the impact of CNFs on amphibians; therefore, it broadened our understanding about ecotoxicological risks associated with their dispersion in freshwater ecosystems and possible contribution to the decline in the populations of anurofauna species.

## 1. INTRODUCTION

The recent scientific and technological development, and the invention of nanomaterials have allowed the creation and production of highly promising and advantageous materials that have been applied to address several challenges associated with conventional Science (Bhagyaraj & Oluwafemi, 2018). Nanomaterials are gaining more and more interest given their unique properties and potential use in a wide range of technological applications. Recent studies have gathered vast information on the use of these materials by the food (Chaudhary et al., 2020; Shafiq et al., 2020), cosmetics (Fytianos et al., 2020; Singh et al., 2020) and civil construction sectors (Firoozi et al., 2020; Singh, 2020), as well as in the manufacture of personal care (Keller et al., 2014; Kaul et al., 2018; Aziz et al., 2019), electronic (Zeb et al., 2019), medicinal and pharmaceutical (Velu et al., 2020; Das et al., 2020; Siddique & Chow, 2020; Kumar et al., 2020) and industrial products (Thomas et al., 2019; Palit & Hussain, 2020), and in different environmental sciences fields (Taran et al., 2020).

Carbon nanofibers (CNFs) that have conductivity and stability similar to that of carbon nanotubes (CNTs) (Lake &Lake, 2014; Mohamed et al., 2019; Yadav et al., 2020) are among the most prominent nanomaterials in recent years. The main features of CNFs distinguishing them from CNTs is the stacking of graphene sheets at different shapes. These sheets produce more edge sites on the outer wall of CNFs than CNTs, and it makes the electron transfer of electroactive analytes easier (Yadav et al., 2020). However, CNFs’ application has mainly focused on catalyst supports (Din et al., 2020), gas-storage systems (Conte et al., 2020), polymer reinforcements (Abdo et al., 2020), probe tips (Cui et al., 2004; Goto et al., 2014) and biosensor development, due to their unique physical and chemical properties (good electrical conductivity, high surface area, biocompatibility, inherent and induced chemical functionalities, and easy manufacture) (Saunier et al., 2020; Senthamizhan et al., 2020).

However, the assessment of ecological risks remain an incipient field involving CNFs, despite their dispersion and distribution in ecosystems - studies carried out with CNTs are much more numerous and comprehensive (Freixa et al., 2018; Gomes et al., 2021). Few investigations with CNFs include assays (Magrez et al., 2006; Brown et al., 2007; Jensen et al., 2012; Kalman et al., 2019) or experiments *in vitro* with invertebrates (Lee et al., 2015) or mammals (DeLorme et al., 2012; Jensen et al., 2012; Warheit, 2019). A small portion of studies *in vivo* has evaluated the effects of these nanomaterials on aquatic freshwater organisms (Chaika et al., 2020; Gomes et al., 2021; Montalvão et al., 2021). However, there is still an important gap in assessments on risk factors posed to, and physiological changes induced by, these compounds in aquatic organisms. Chaika et al. (2020) assessed CNF effects on the digestive system of different freshwater invertebrates (Families: Gammaridae, Ephemerellidae and Chironomidae), but they did not observe any histopathotoxic effect on animals’ gastrointestinal tract. In fact, these authors have shown the ability of Gammarus suifunensis to biodegrade CNFs (Chaika et al. 2020). Gomes et al. (2021) have evidenced that CNFs can be transferred by an experimental food chain (*Eiseia* fetida > *Danio rerio > Oreochromis niloticus*) and cause mutagenic and cytotoxic damage at the uppermost trophic level. Montalvão et al. (2021) reported that dragonfly larvae (*Aphylla williamsoni*) short-term exposure (48 h) to CNFs induced predictive changes in REDOX imbalance and neurotoxicity - this finding was inferred by suppressing the activity of acetylcholinesterase (AChE).

Therefore, the inconclusive character of the investigative scenario about CNFs’ toxicity, as well as the gaps on knowledge about the impact of these nanomaterials on several groups of invertebrates and vertebrates are clear factors, so far. Amphibians are among these groups, but, although they have priceless ecological importance (Hocking & Babbitt, 2014), they have never been the subject of investigations involving CNFs. Our knowledge about the toxicity of carbon-based nanomaterials (CNs) in amphibians is restricted to information available in reports by Saria et al. (2014) and Zhao et al. (2020). These authors were the first to show that the short-term exposure of *Xenopus laevis* tadpoles to multi-walled carbon nanotubes (MWCNTs) induced oxidative stress and caused damage to animals’ erythrocyte DNA. Zhao et al. (2020) reported MWCNT accumulation in different organs of tadpoles belonging to species X. *tropicalis* increased their lethality rate and changed their heart rate. Thus, it is imperative carrying out further studies to assess how CNTs can have impact on the anurofauna and ecotoxicological effects of CNFs. These complementary investigations are essential, since amphibians are organisms sensitive to changes in their habitats (Roy, 2002; Wagner et al., 2014; Rohman et al., 2020), and are included in the list of animals presenting significant population decline in recent years (Green et al., 2020).

Accordingly, we evaluated the likely toxicological effects of CNFs on tadpoles belonging to neotropical species *Physalaemus cuvieri* (Anura, Leptodactylidae). This species is exclusively distributed in South America and is typical of open biomes, such as Cerrado, Caatinga, Chaco and Llanos (Mijares et al., 2011; De-Oliveira-Miranda et al., 2019). Although the species is currently categorized as of “little concern” by the International Union for Conservation of Nature (stable, least concern, version 2020-3) (IUCN, 2020), its wide geographical distribution and presumed large populations, are features turning them into interesting model systems, since they can inhabit freshwater environments subjected to different pollution types, including CNFs. From different biomarkers, We herein aimed at testing the hypothesis that short exposure to CNFs (at environmentally relevant concentrations) induces changes in the nutritional status, metabolic changes altering REDOX homeostasis into oxidative stress, and cytotoxic and neurotoxic changes in these animals. To the best of our knowledge, this is the first report on the biological impact of CNFs on a specific amphibian species. Therefore, this study has broadened our understanding about ecological risks associated with water pollution by these nanomaterials, as well as motivated further investigations on the impact of CNs on amphibians’ health and on the dynamics of their natural populations.

## 2. MATERIALS AND METHODS

### 2.1. Carbon nanofibers

We used pyrolytically stripped CNFs (i.e., polyaromatic hydrocarbons removed from fibers’ surface) provided by Sigma-Aldrich (San Luis, Missouri, USA) - their detailed chemical featuring was presented by Gomes et al. (2021). These pollutants are a mix of different sized and shaped CNFs [from 60 to 100 nm (mean: 86.85 ± 1.80 nm)], including the ones with open and clearly curved tips (Figure 1). According to the manufacturer, and as seen in the photoelectric micrographs taken during the transmission electron microscopy analysis, the assessed CNFs have different metallic particles (Ca, Si, S, Na, Mg and Fe), used as catalysts (Figure 1).

**Figure 1.**
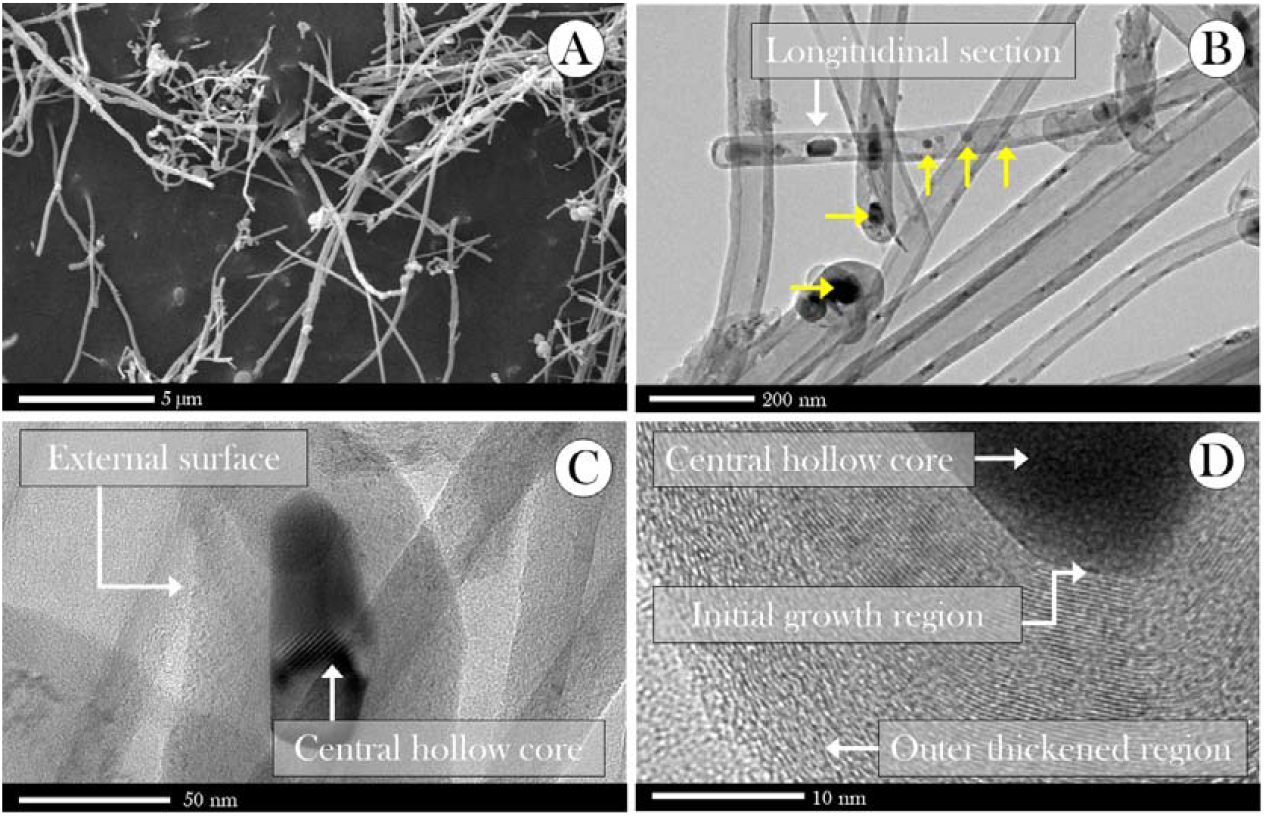
(A) Scanning electron microscopy images and (B-D) transmission electron microscopy images of a CNF film, at different magnifications. Yellow arrows point out the presence of metallic particles inside CNFs, or around their surface, as shown in (D).

### 2.2. Model system and experimental design

We used tadpoles belonging to species *P. cuvieri* (Anura, Leptodactylidae) as model system to assess the aquatic toxicity of CNFs. Its wide geographical distribution in South America (Miranda et al., 2019), stability and population abundance in the occupied areas (Frost, 2017), good adaptability to laboratory environment, early biological response to changes in its environment, and use in recent (eco) toxicological studies justify their choice as model in our study (Herek et al., 2020; Araújo et al., 2020ab; Rutkoski et al., 2020). All tadpoles used in the experiment came from an ovigerous mass with approximately 1,500 eggs, based on Pupin et al. (2010). The ovigerous mass was collected in a lentic environment (Urutaí, GO, Brazil) surrounded by native Cerrado biome, under license n. 73339-1 - issued by the Brazilian Biodiversity Authorization and Information System (SISBIO/MMA/ICMBio).

Eggs were kept in aquarium (40.1 x 45.3 x 63.5 cm) filled with 80 L of naturally dechlorinated water (for at least 24 h), under controlled 12h light-dark photoperiod and temperature (26 °C ± 1 ° C - similar to that of the natural environment) conditions, and constant aeration (by air compressors) from the time they were taken to the laboratory. Animals were fed once a day (ad libitum) with commercial fish food (formula: 45% crude protein, 14% ether extract, 5% crude fiber, 14% mineral matter and 87% dry matter). Tadpoles remained under the aforementioned conditions until they reached stage 27G (body biomass: 70 mg ± 4.1 mg; and total length: 20.1 mm ± 0.7 mm - mean ± SEM)., after egg hatching, based on Gosner (1960). The healthy tadpoles (i.e., the ones presenting normal swimming movements and no morphological deformities or apparent lesions) were divided into three experimental groups (n = 195 tadpoles/each - 13 replicates composed of 15 animals/each). The control group (C) was composed of tadpoles kept in dechlorinate tap water (CNFs free) and groups CNF-I and CNF-II comprised animals exposed to water added with CNFs at concentrations of 1 and 10 mg/L, respectively (see below).

### 2.3. Exposure conditions and CNF concentrations

All experimental groups were kept in polyethylene containers filled with 180 mL of dechlorinated water where CNFs were diluted in. Exposure time was set at 48 h (static system) to simulate ephemeral exposure. Animals’ food was kept during exposure - commercial feed was offered once a day. Concentrations of the tested CNFs were defined based on aquatic CNT concentrations, due to lack of information about environmental concentrations recorded for CNFs. Therefore, previous studies evaluating CNTs’ toxicity in different experimental models based on concentrations ranging from 0.1 to 100 mg/L were taken as basis to select CNF concentrations used in the current research (Mouchet et al., 2007; Mouchet et al., 2009; Mouchet et al., 2010; Mouchet et al., 2011; Bourdiol et al., 2013; Saria et al., 2014; Verneuil et al., 2015; Zhao et al., 2020; Tavabe et al., 2020). We herein applied the monitoring MWCNT data recorded by Nezhadheydari et al. (2019) in aquatic environments and the experimental design proposed by Tavabe et al. (2020). The aforementioned authors observed concentration up to 20 mg/L of these materials, and it proved the significant changes in it (ng/L to mg/L). Concentrations tested in the current study were environmentally relevant, and it takes the present experimental design closer to realistic CNF-pollution conditions. The herein adopted concentrations represented both optimistic (less pollution; 1 mg/L) and pessimistic (higher pollution; 10 mg/L) conditions.

### 2.4. Toxicity biomarkers

#### 2.4.1. Biochemical assessments

##### 2.4.1.1. Sample preparation

Samples were prepared based on Guimarães et al. (2021), with modifications, to evaluate the biochemical parameters. In total, 144 tadpoles were used per experimental group (n = 12 samples, composed of a pool of 12 animals/each). These animals were weighed, macerated in 1 mL of phosphate buffered saline (PBS) solution and centrifuged at 13,000 rpm, for 5 min (at 4°C). The supernatant was separated into aliquots to be used in different biochemical evaluations. Whole bodies were used due to technical limitations in isolating certain organs from small animals. Unlike assessments in adult specimens, organ-specific biochemical assessment carried out in tadpoles require highly accurate dissection due to their small sized-bodies, which makes it difficult processing large numbers of samples under time constraint (Khan et al. 2015). Organ “contamination” by organic matter and/or by other particles consumed by tadpoles can be a bias for the biochemical analysis applied to organs during dissection time (Lusher et al. 2017; Guimarães et al., 2021).

##### 2.4.1.2. Nutritional status

Different pollutants can affect the nutritional status of tadpoles (Bharatraj & Yathapu, 2018); therefore, we evaluated total soluble carbohydrate, total protein, and triglyceride concentrations in different tissues of the exposed animals. Total soluble carbohydrate levels were determined through the Dubois method (Dubois et al., 1956) - detailed by Estrela et al. (2021). Protein level was determined in commercial kit (Bioténica Ind. Com. LTD, Varginha, MG, Brazil. CAS number: 10.009.00), based on biuret reaction (Gornall et al., 1949; Henry et al., 1957). Triglyceride levels were evaluated based on Bucolo & Davis (1973) by using a commercial kit (Bioténica Ind. Com. LTD, Varginha, MG, Brazil. CAS number: 10.010.00).

##### 2.4.1.3. REDOX state

###### 2.4.1.3.1. Oxidative stress biomarkers

Likely oxidative stress increase was assessed based on indirect nitric oxide (NO) determination, REDOX regulated processes through nitrite measurement (Soneja et al. 2005), thiobarbituric acid reactive species (TBARS) [predictive of lipid peroxidation (De-Leon & Borges, 2020)], reactive oxygen species (ROS) production and on hydrogen peroxide (H_2_O_2_) - which plays essential role in responses to oxidative stress, in different cell types (Sies, 2020; Sies et al., 2020). The Griess colorimetric reaction [based on Bryan et al., (2007)] was used to measure nitrite concentrations. TBARS levels were determined based on procedures described by Ohkawa et al. (1979) and modified by Sachett et al. (2020). H_2_O_2_ and ROS production was assessed based on the methodological procedures proposed by Elnemma et al. (2004) and Maharajan et al. (2018), respectively.

###### 2.4.1.3.2. Antioxidant response biomarkers

The activation or suppression of antioxidant activity in animals exposed to different CNF concentrations was evaluated by determining catalase and superoxide dismutase (SOD) activity. These enzymes are considered first-line antioxidants important for defense strategies against oxidative stress (Ighodaro & Akinloye, 2018). Catalase activity was assessed based on Sinha et al. (1972) [see details in Montalvão et al. (2021)] and SOD was determined according to the method originally described by Del-Maestro & McDonald (1985) and adapted by Estrela et al. (2021).

##### 2.4.1.4. Cytotoxicity

Blood samples were collected to assess cytotoxic effects induced by CNFs through erythrocytic apoptosis or necrosis. Procedures like those described by Singla &Dhawan (2013) and García-Rodríguez et al. (2013) were herein adopted. Briefly, 0.5-1.0 *μL* of blood from two animals in each group (n= 16 por grupo) was mixed to 200 *μL* of PBS. Subsequently, 50 *μL* of acridine orange dye solution (AO) and 50 *μL* of ethidium bromide (EB) solution (both at 1 μg/mL) were added to the mix, which was incubated at room temperature, for 5 min. Samples were then centrifuged (at 13,000 rpm and 4°C, for 5 min). The pellet was resuspended, placed on slide and covered with a glass cover slip after the supernatant was discarded. A barrier filter for immediate evaluation under fluorescence microscope (BEL Engineering^®^, model FLUO3 - excitation 510-560 nm) was used in the experiment. The total number of 100 cells from each slide was scored for apoptosis extent quantification. Living cells were green, apoptotic cells were orange and presented fragmented nuclei, and necrotic cells were red (Kasibhatla et al., 2006; Singla &Dhawan, 2013). The rate of each cell type, in each animal, was calculated.

##### 2.4.1.4. Neurotoxicity

The induction of likely neurotoxic effect caused by CNFs was evaluated by determining acetylcholinesterase (AChE) activity based on the method by Ellman et al. (1961) and the activity of butyrylcholinesterase (BChE) - also known as serum cholinesterase or pseudocholinesterase – based on Silva et al. (2020). We also evaluated whether CNFs could change the viability of neuromats living on tadpoles’ surface (Russell, 1976) – this feature has been considered a good ecotoxicological biomarker (Guimarães et al., 2021). Accordingly, 10 living tadpoles from each group were exposed (for 15 min) to water reconstituted with 4- (4-Diethylaminostyryl) −1-methylpyridinium iodide (4-Di-2-ASP) at 5 mM, similar to procedures adopted by Krupa et al. (2020) and Guimarães et al. (2021). Subsequently, animals were anesthetized (on ice) and taken to fluorescence microscope (BEL Engineering^®^, model FLUO3 - excitation 510-560 nm) to have images of their heads and tails captured. The number of neuromats was manually determined; neuromats located on the sides of the tadpoles were excluded because they were out of focus or absent, due to their position in the microscope. We also excluded the lower part of their head and their back-posterior region, which overall had expressive amounts of non-specific coloring. Neuromats on the head and tail sides were quantified, as shown in Figure 2.

**Figure 2.**
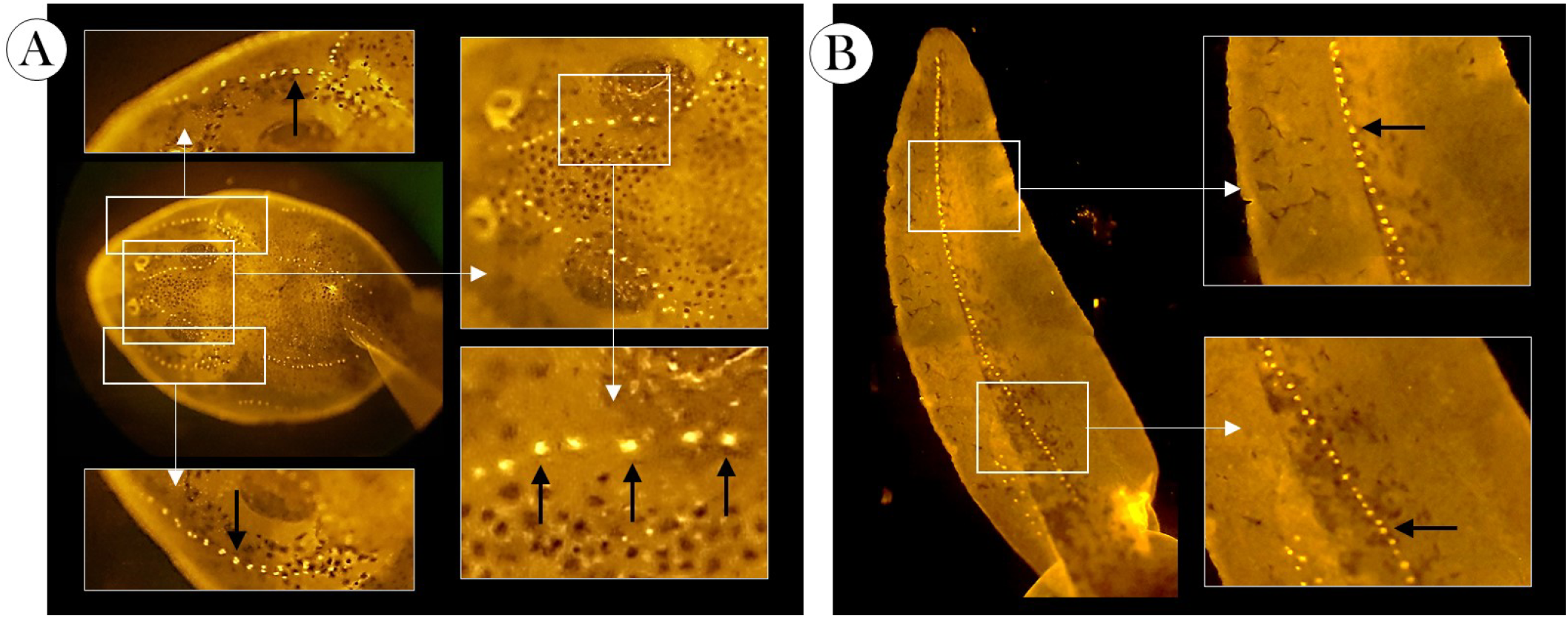
Representative images of the (A) head and (B) tail regions of *P. cuvieri* tadpoles, whose number of superficial neuromats were counted.

### 2.6. CNF accumulation

CNF accumulation was estimated by determining total organic carbon (TOC) concentrations by taking into consideration the specific quantification of CNs in environmental and biological samples. This process is a huge challenge, given the lack of accessible standard methods to quantify these nanomaterials (Wang et al., 2013; Chang et al., 2014; Bourdiol et al., 2015; Petersen et al., 2016). We herein adopted the Walkley-Black method used by Schwab et al. (2011) and Gomes et al. (2021); this method is based on using dichromate as oxidizer in acid medium (Walkley & Black, 1934). Detailed methodological procedures can be observed in a previous study carried out by our research team (Gomes et al., 2021). Results were expressed in “g of TOC/kg of body biomass” (n = 16/group, 8 samples composed of a pool of 2 animals/each).

### 2.7. Visual assessment

Nine tadpoles randomly selected from each group were euthanized on ice and incubated in acridine orange dye (AO) and ethidium bromide (EB) solution (both at 1 μg/mL), at room temperature for 10 min, in addition to CNF accumulation estimates. Such procedure allowed better differentiating different regions of animal’s body displaying accumulated CNFs. Their animals were captured in fluorescence microscope (BEL Engineering^®^, model FLUO3 - excitation 510-560 nm) for further qualitative evaluation.

### 2.8. Statistical analysis

GraphPad Prism Software Version 8.0 (San Diego, CA, USA) was used to the statistical analyses. Initially, data were checked for normality and homogeneity variance deviations before the analysis. Normality data were assessed through Shapiro-Wilks test, and variance homogeneity was assessed through Bartlette’s test. Multiple comparisons were performed by applying one-way ANOVA and Tukey’s post-hoc analysis to non-parametric data or Kruskal-Wallis test, Dunn’s post-hoc test to non-parametric data. Significance levels were set at Type I error (p) values lower than 0.05, 0.01 or 0.001.

## 3. RESULTS

By assuming the possible interference of CNFs in tadpoles’ energy metabolism, we evaluated the concentration of different macromolecules. Both concentrations recorded for the tested CNFs have significantly reduced total soluble carbohydrate and total protein levels in these animals, except for triglyceride levels, whose reduction was only observed in animals in the CNF-II group (Figure 3) - with no concentration-response effect. Based on our data, CNFs induced nitrite production increase in animals belonging to group CNF-I (Figure 4A), as well as in H_2_O_2_ and ROS production in both groups exposed to nanomaterials (Figure 4B-C, respectively), and TBARS production in animals kept in water added with 10 mg/L of CNFs (Figure 4D).

**Figure 3.**
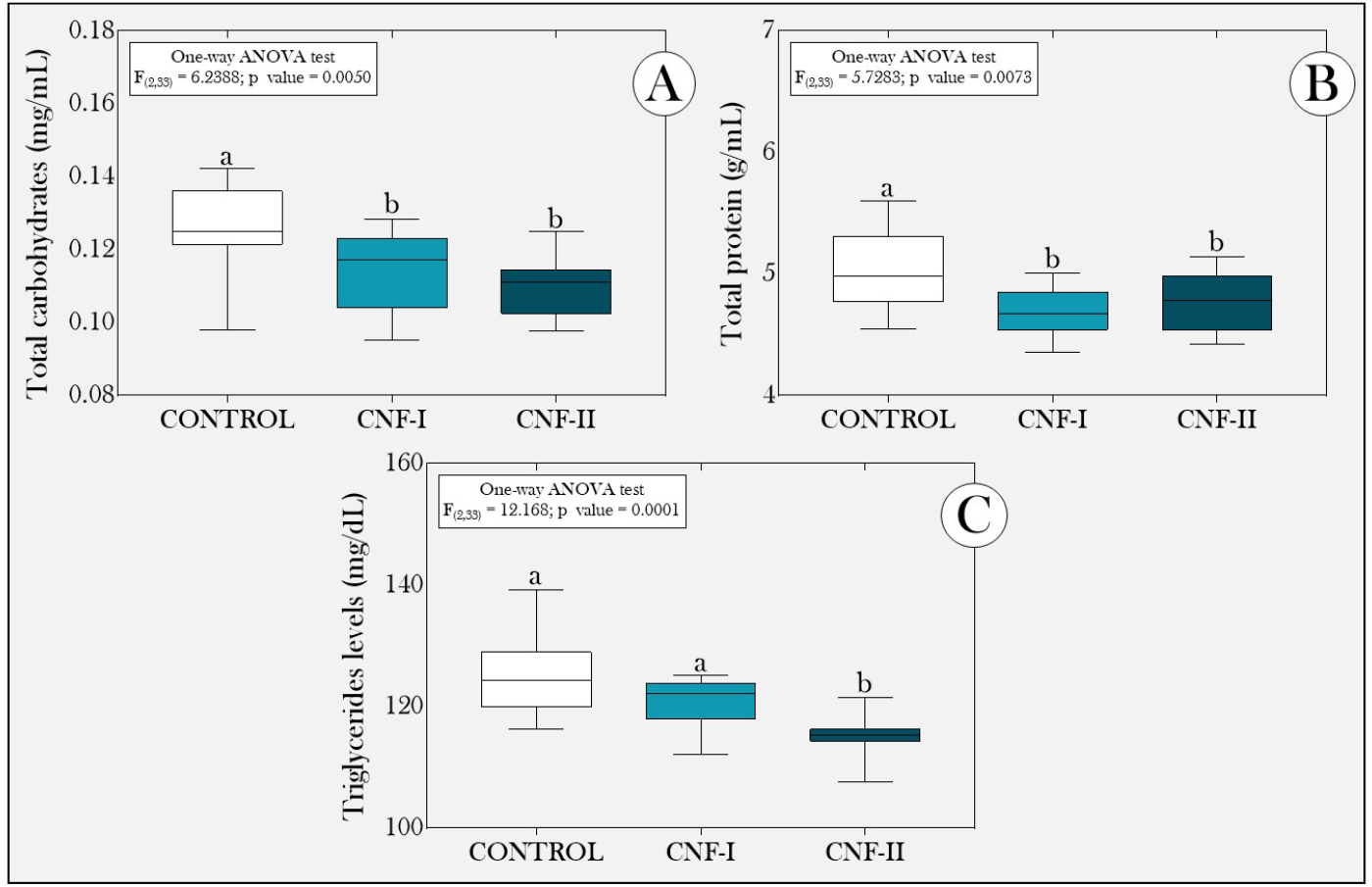
Boxplot of (A) total soluble carbohydrate, (B) total protein and (C) triglycerides concentration in *P. cuvieri* tadpoles exposed, or not, to different CNF concentrations. Summaries of statistical analyses are shown in the upper left corner of the figures. Different lowercase letters indicate significant differences between experimental groups. CONTROL: group of tadpoles not exposed to CNFs. CNF-I and CNF-II groups: tadpoles exposed to carbon nanofibers at concentrations of 1 and 10 mg/L, respectively, n = 144 tadpoles/group, 12 samples/group composed of a pool of 12 animals/each.

**Figure 4.**
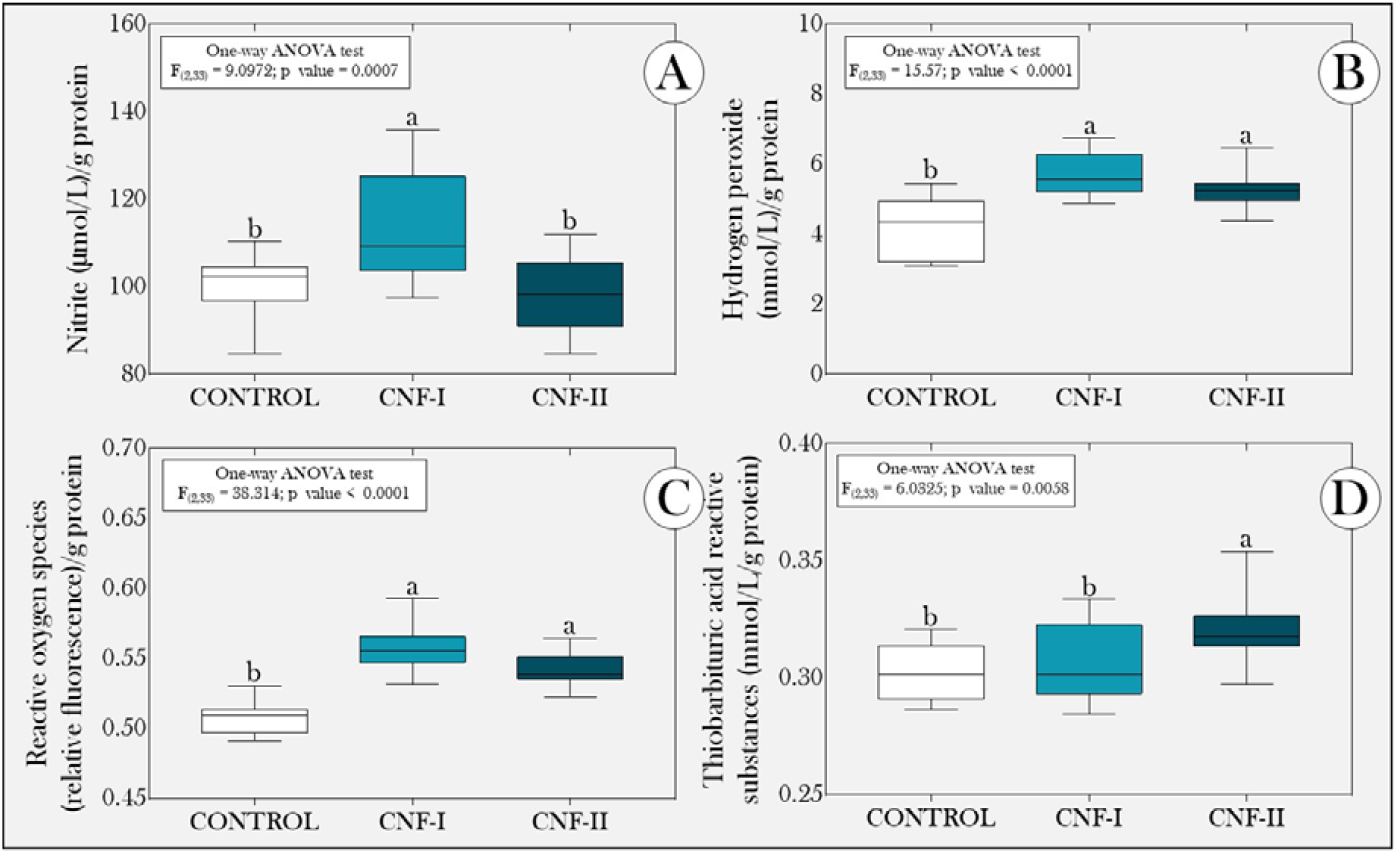
Boxplot of (A) nitrite, (B) hydrogen peroxide, (C) reactive oxygen species (ROS) and (D) thiobarbituric acid reactive substances concentrations in *P. cuvieri* tadpoles exposed, or not, to different CNF concentrations. Summaries of statistical analyses are shown in the upper left corner of the figures. Different lowercase letters indicate significant differences between experimental groups. CONTROL group: tadpoles not exposed to CNFs. CNF-I and CNF-II groups: tadpoles exposed to carbon nanofibers at concentrations of 1 and 10 mg / L, respectively (n = 144 tadpoles/group, 12 samples/group composed of a pool of 12 animals/each).

We observed significant increase in SOD and catalase levels in animals exposed to CNFs, but no concentration-response effect (Figure 5A-B, respectively). SOD levels were positively and significantly correlated to H_2_O_2_, ROS and TBARS levels (Table 1). Catalase concentrations were correlated to H_2_O_2_ and ROS production (Table 1).

**Figure 5.**
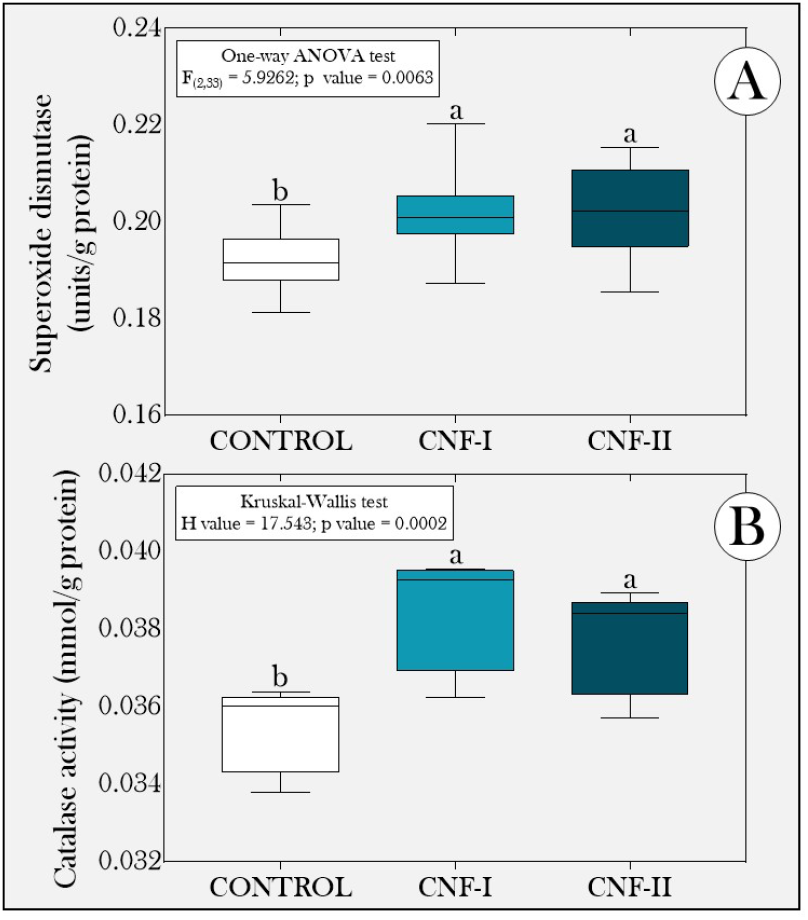
Boxplot of (A) superoxide and (B) dismutase concentration in tadpoles belonging to species *P. cuvieri* exposed, or not, to different CNF concentrations. Summaries of statistical analyses are shown in the upper left corner of the figures. Different lowercase letters indicate significant differences between experimental groups. CONTROL group: tadpoles not exposed to CNFs. CNF-I and CNF-II groups: tadpoles exposed to carbon nanofibers at concentrations of 1 and 10 mg/L, respectively (n = 144 tadpoles/group, 12 samples/group composed of a pool of 12 animals/each).

**Table 1.**
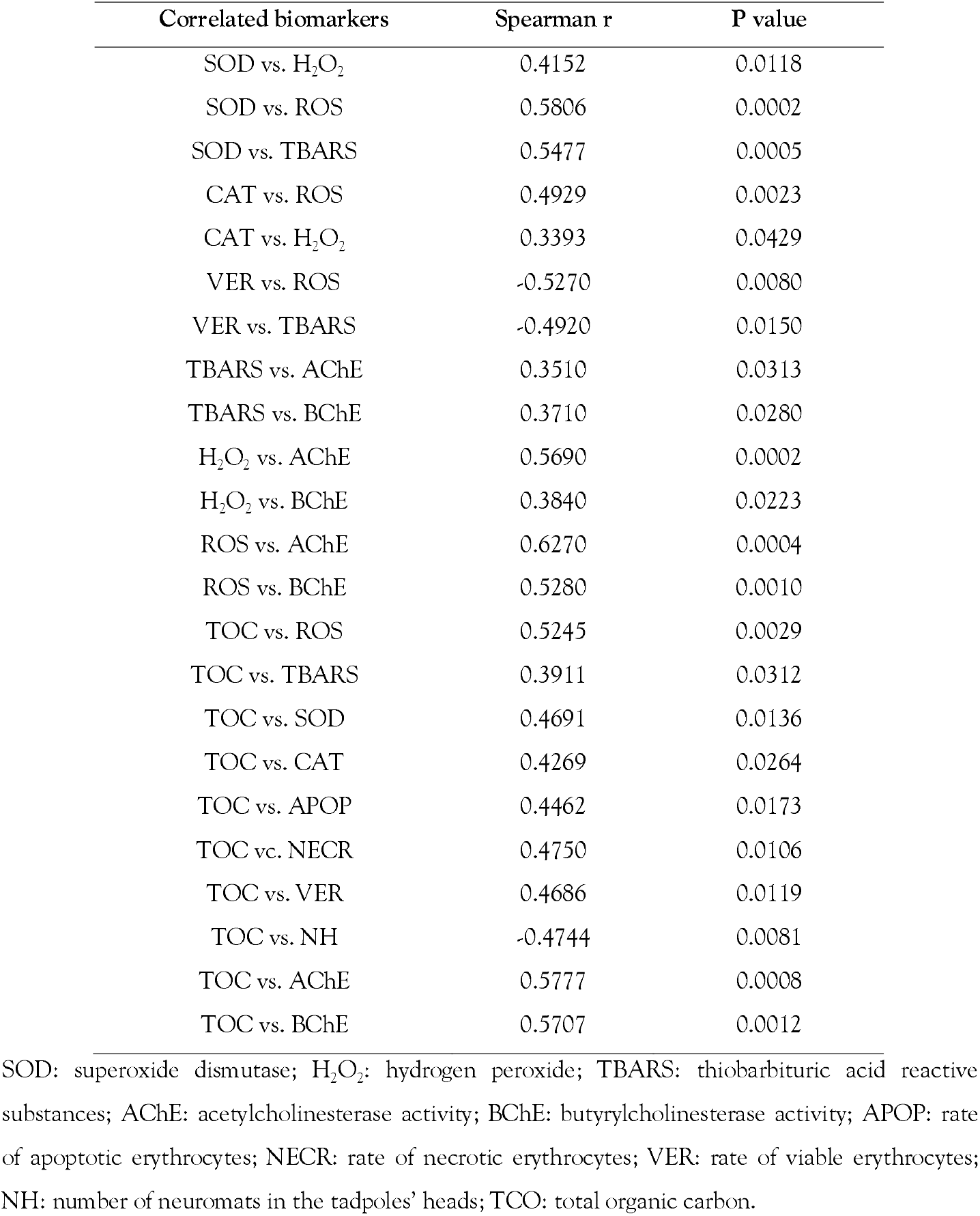
Summary of correlation analyses carried out between different biochemical biomarkers.

Based on our data, there was cytotoxic effect induced by CNFs on tadpoles’ erythrocytes. Figure 6 depicts that the groups exposed to nanomaterials recorded lower rates of viable erythrocytes and, consequently, higher rates of apoptotic and necrotic cells than animals in the control group. The rate of viable cells was negatively and significantly correlated to ROS and TBARS concentrations (Table 1). According to the neurotoxic evaluation, there was AChE and BChE increase in animals exposed to nanomaterials, and this finding suggests the stimulatory effect induced by CNFs on tadpoles’ cholinergic system (Figure 7A-B, respectively). On the other hand, tadpoles exposed to CNFs showed smaller number of superficial neuromats in their heads (Figure 8A) and a larger amount of them in their tail (Figure 8B), but no concentration-response effect. However, the total number of neuromats (head + tail) did not differ between experimental groups (Figure 8C).

**Figure 6.**
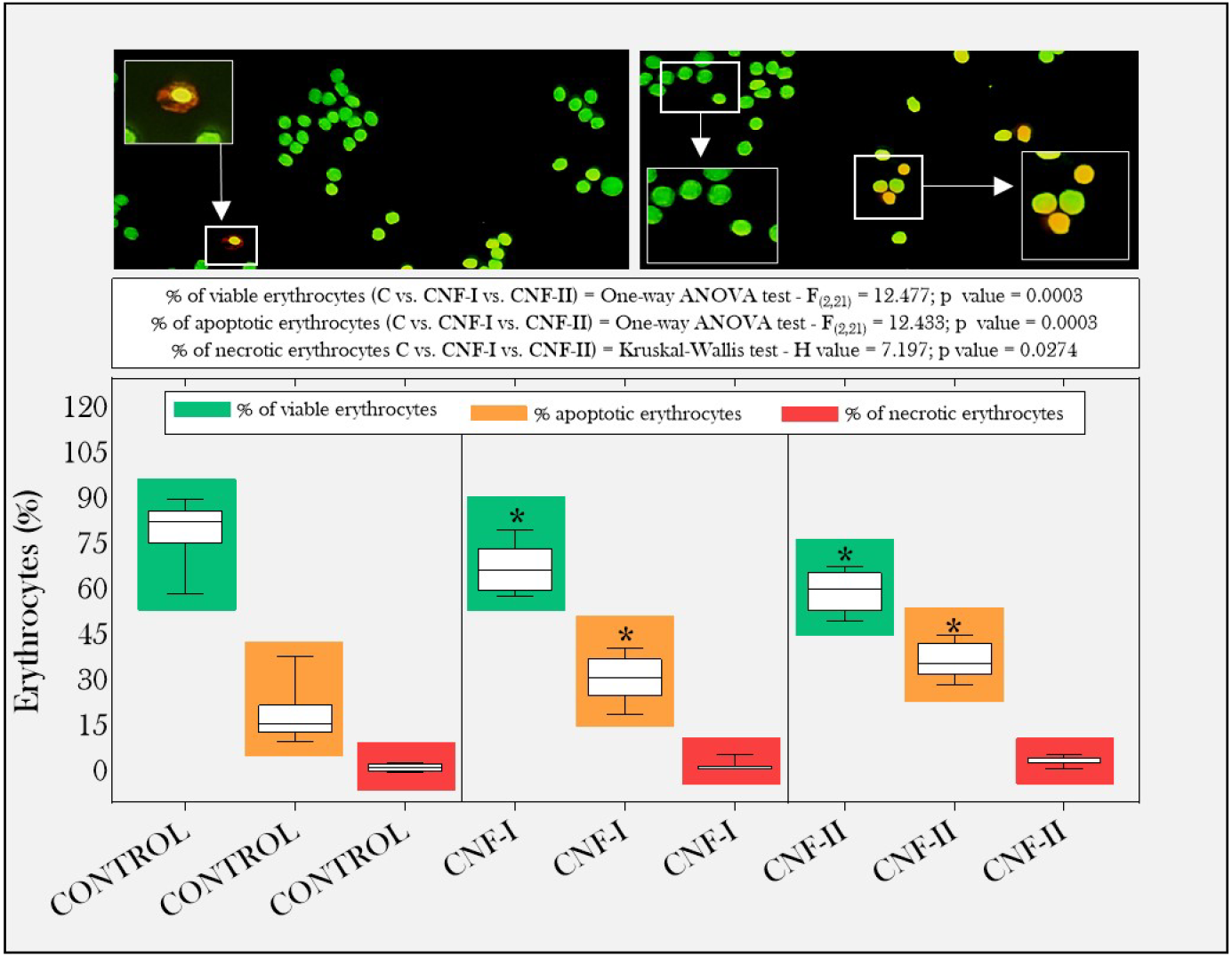
Boxplot of viable (green), apoptotic (orange) and necrotic (red) erythrocyte rates in *P. cuvieri* tadpoles exposed, or not, to different CNF concentrations. Fluorescence images representative of acridine orange and ethidium bromide staining are presented above the boxplot. Summaries of statistical analyses are shown in the upper left corner of the figures. Asterisks indicate differences between the respective cell types from each group exposed to CNFs and from the control group. CONTROL group: tadpoles not exposed to CNFs. CNF-I and CNF-II groups: tadpoles exposed to carbon nanofibers at concentrations of 1 and 10 mg/L, respectively (n = 16 tadpoles/group, 8 samples, composed of a pool of two animals/each).

**Figure 7.**
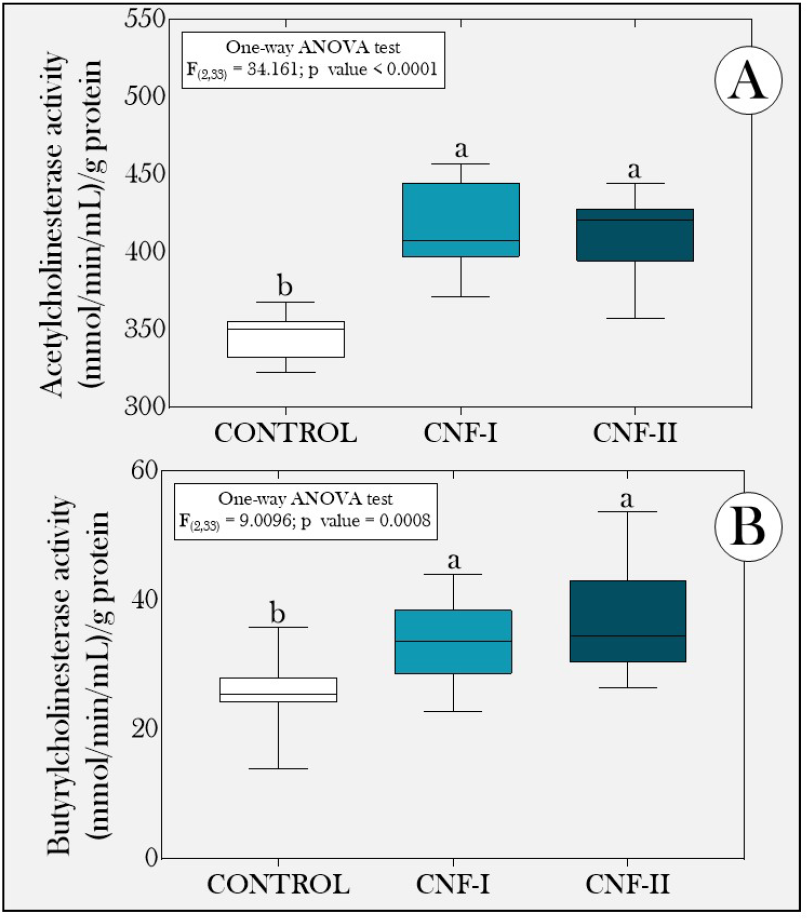
Boxplot of (A) acetylcholinesterase and (B) butyrylcholinesterase activity in *P. cuvieri* tadpoles exposed, or not, to different CNF concentrations. Summaries of statistical analyses are shown in the upper left corner of the figures. Different lowercase letters indicate significant differences between experimental groups. CONTROL group: tadpoles not exposed to CNFs. CNF-I and CNF-II groups: tadpoles exposed to carbon nanofibers at concentrations of 1 and 10 mg/L, respectively (n = 144 tadpoles/group, 12 samples/group composed of a pool of 12 animals/each).

**Figure 8.**
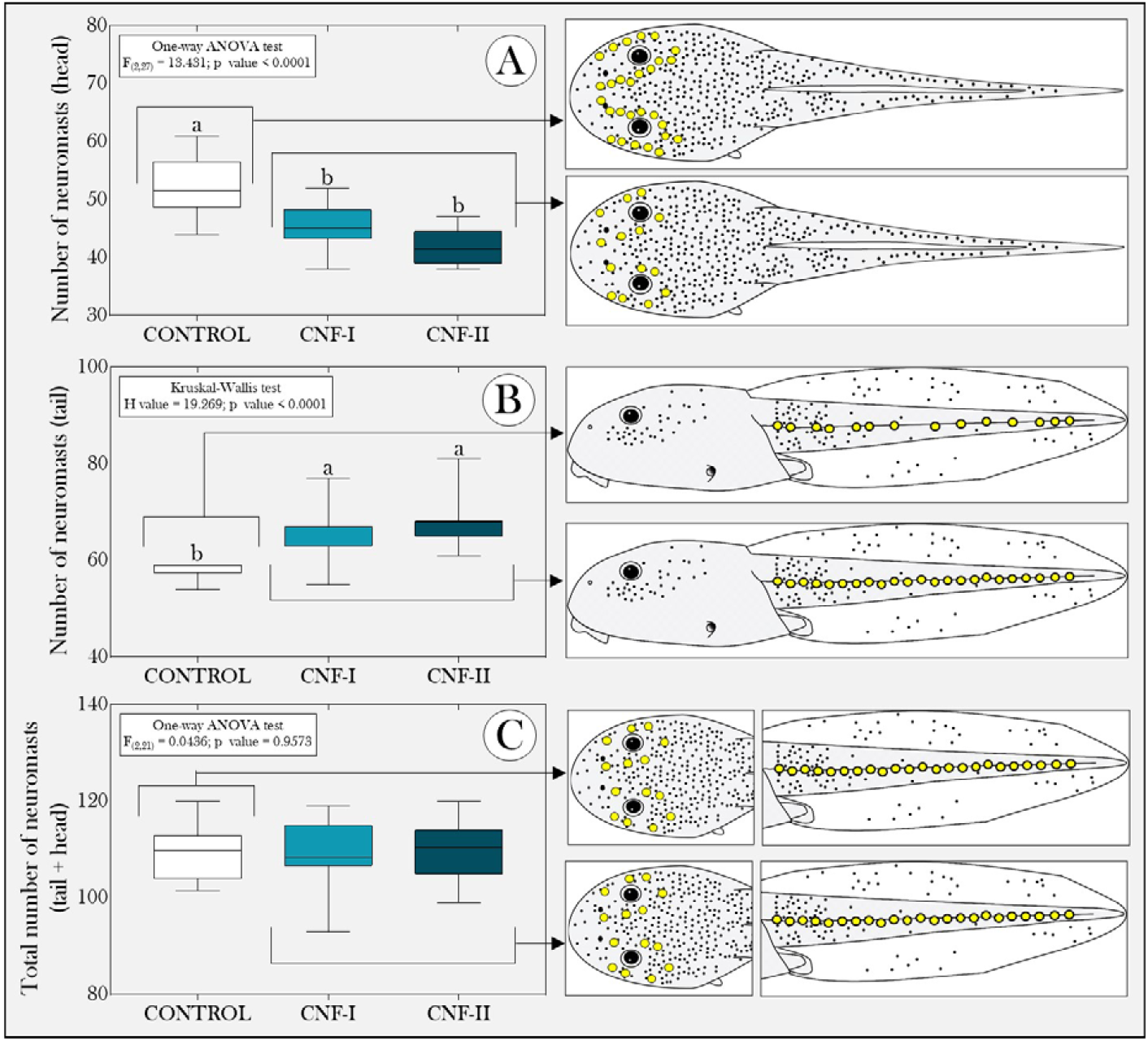
Boxplot of number of superficial neuromats in the (A) head and (B) tail of tadpoles, and the total number (C) of *P. cuvieri* tadpoles exposed, or not, to different CNF concentrations. Summaries of statistical analyses are shown in the upper left corner of the figures. Different lowercase letters indicate significant differences between experimental groups. CONTROL group: tadpoles not exposed to CNFs. CNF-I and CNF-II groups: tadpoles exposed to carbon nanofibers at concentrations of 1 and 10 mg/L, respectively (n = 10 tadpoles/group).

Finally, we observed the accumulation of nanomaterials in tadpoles belonging to groups CNF-I and CNF-II - this finding was inferred based on TOC concentrations (Figure 10A) and on animals’ visual evaluation (Figure 10B-C) - with concentration-response effect. We noticed significant CNF accumulation in animals’ gastrointestinal tract; it prevailed in the ones exposed to the highest CNF concentration (10 mg/L). These data have confirmed that CNFs were ingested by tadpoles; the statistical analyses have shown significant correlation among the accumulation of these nanomaterials, different biomarkers predictive of oxidative stress (ROS and TBARS), antioxidant activity (SOD and CAT), as well as cytotoxic (viable, apoptotic, and necrotic erythrocytes) and neurotoxic effect (number of neuroblasts in tadpoles, as well as AChE and BChE activity in these models) (Table 1).

**Figure 10.**
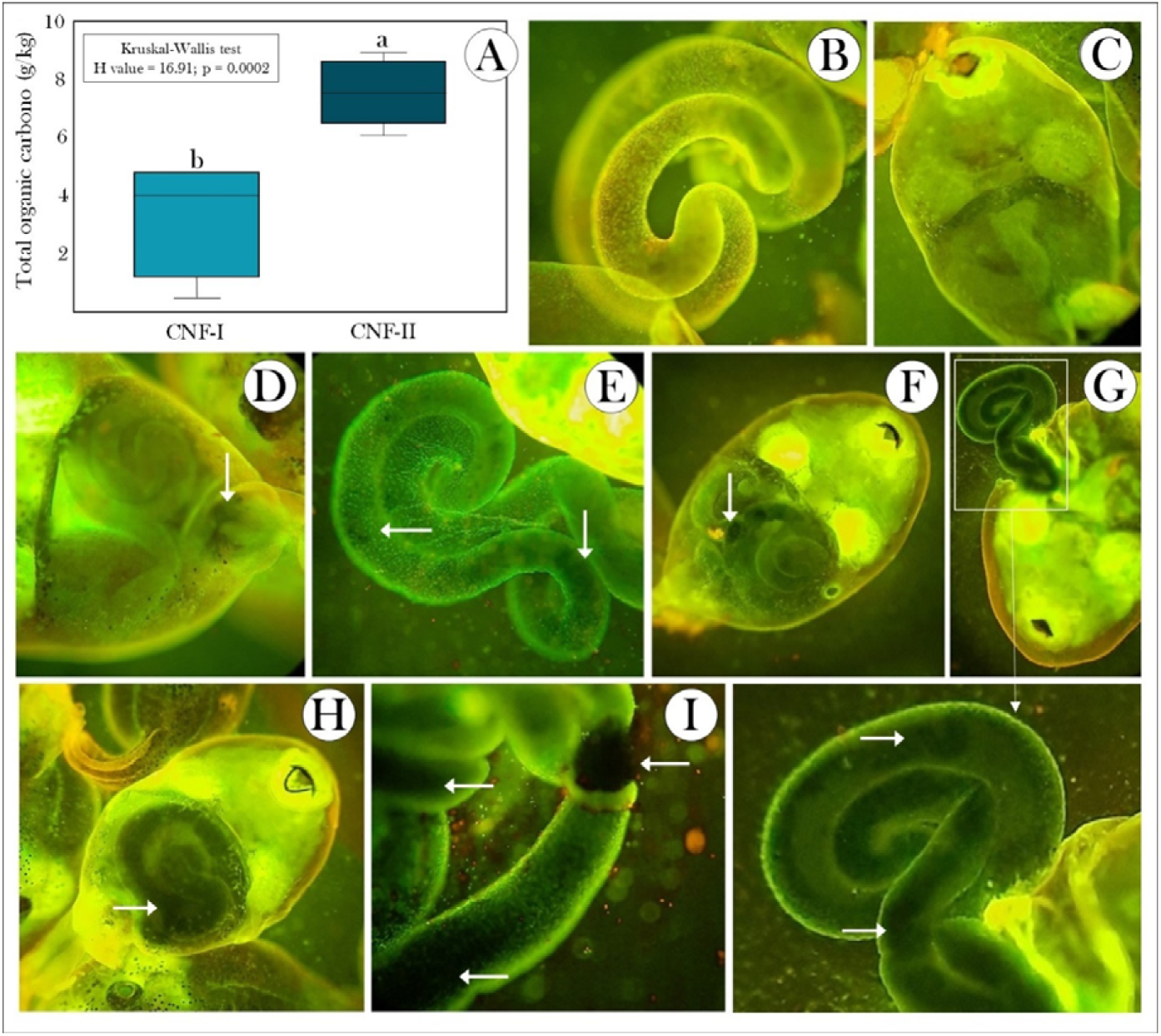
Boxplot of (A) total organic carbon concentration and (B-I) representative images of *P. cuvieri* tadpoles exposed, or not, to different CNF concentrations. “A”, the summary of the statistical analysis is shown in the upper left corner of the figure. Different lowercase letters indicate significant differences between experimental groups. Background TOC concentrations in tadpoles in the control group were detected and subtracted from that of CNFs-exposed samples. (B-C): representative images of *P. cuvieri* tadpoles not exposed to CNFs, (D-F) images of animals exposed to the lowest (1 mg / L) and (G-I) highest concentrations (10 mg / L) of the pollutant. CNF-I and CNF-II groups: tadpoles exposed to carbon nanofibers at concentrations of 1 and 10 mg/L, respectively. Representative images of n = 9 tadpoles/group. “A”, data about n = 16/group, 8 samples, composed of a pool of two animals/each. Arrows indicate CNF accumulation in animals.

## 4. DISCUSSION

Identifying and featuring the effect of organisms’ exposure to different pollutants and contaminants are essential procedures to assess ecotoxicological risks associated with environmental pollution (Eason & O’Halloran, 2002; O’Halloran, 2006; Rand, 2020). Furthermore, the aforementioned studies can generate important subsidies for further remediation/mitigation measures to prevent the occurrence of lethal effects on the herein assessed animals (Madear et al., 2020; Wang et al., 2020). Accordingly, our study was the first to show how CNFs can have impact on the health of anurofauna. This finding reinforces the toxicological potential of these nanomaterials inferred through tests carried out *in vitro* and *in vivo* with organisms from other taxonomic groups (Magrez et al., 2006; Brown et al., 2007; Jensen et al., 2012; DeLorme et al., 2012; Jensen et al., 2012; Lee et al., 2015; Kalman et al., 2019; Warheit, 2019).

The herein evidenced significant reduction in total soluble carbohydrate, total protein and triglyceride concentrations in *P. cuvieri* tadpoles exposed to CNFs (Figure 3) and to other types of pollutants (e.g.: atrazine: Dornelles & Oliveira, 2014; glyphosate: Dornelles & Oliveira, 2016; quinclorac: Dornelles & Oliveira, 2014; basudin: Ezemonye & Ilechie, 2007; naphthenic acids: Melvin et al., 2013; polycyclic aromatic hydrocarbons: Gendron et al., 1994, among others) has proven that CNFs can have impact on the energy metabolism of these models. Therefore, our results have suggested the direct or indirect activation of metabolic pathways related to glycogenolysis, proteolysis and lipolysis by CNFs. In this case, assumingly, the high energy consumption demanded by physiological processes of antioxidant defense, whose association was previously discussed (Strong et al., 2017) can explain the lower concentrations of the assessed macromolecules. Therefore, it is possible stating that CNF accumulation in animals’ gastrointestinal system (Figure 9) was affected by nutrient absorption either by saconsumption by tadpolespce occupation in the intestinal lumen [like reports involving microplastic consumption by tadpoles (Araújo et al., 2020a)] or by its negative effects on intestinal absorptive cells or on tadpoles’ hepatic system. We could rule out the hypothesis that CNF accumulation in tadpoles’ gastrointestinal tract itself can trigger physiological mechanisms capable of diminishing these models’ food capture motivation, as also proposed by Araújo & Malafaia (2020). The false satiety feeling of the animals can reduce carbohydrate, lipid, and tissue protein rates in tadpoles’ bodies. In any case, nutritional deficits can have broader ecological consequences in tadpoles, regardless of the physiological mechanisms altered during exposure to CNFs, since the diverting energy from other processes, such as growth and development, to maintain physiological homeostasis, often has negative effect on these animals’ health.

On the other hand, our data have evidenced CNFs’ ability to induce oxidative stress increase, which was inferred based on ROS, H_2_O_2_ and TBARS concentrations (Figure 4) - SOD and catalase activation (Figure 5) was not enough to maintain homeostasis REDOX in tadpoles exposed to the tested nanomaterials. Although the literature about studies involving amphibians’ exposure to any CN carried out *in vivo* is scarce, our data have corroborated results in reports by Sari et al. (2014). These authors reported that increased H_2_O_2_, glutathione reductase, SOD and catalase rates in tadpoles belonging to species *Xenopus leavis* were exposure-time (2, 4, 8, 12 and 24 h) and MWCNT- (0.1, 1 and 10 mg/L) dependent.

From the biochemical viewpoint, the most important enzymatic pathways for antioxidant defense against ROS are those involving SOD, since they convert the superoxide anion radical (O_2_) into H_2_O_2_, and catalase, which converts H_2_O_2_ into FLO molecules and O_2_ (Lee et al., 2018; Ransy et al., 2020; Damiano et al., 2020). Based on such an information, it is tempting to speculate that the increased oxidative stress observed in our study can be explained by different responses to CNFs. One possibility for this statement could be related to the negative effect of CNFs on catalases’ molecular structure, because it decreases catalases’ enzymatic efficiency or influences its affinity with the substrate. It is so because H_2_O_2_ molecules formed through SOD activity would not be neutralized by catalase, although its activity increases in animals exposed to CNFs. In this case, even greater increase would be necessary to balance SOD and catalase activity. However, assumingly, H_2_O_2_ is released as the product from other metabolic routes [see review by Hernandez et al. (2012)], catalyzed by enzymes, such as alcohol (Siebum et al., 2006; Ferreira et al., 2010; Turner, 2011), glucose (Zhou et al., 2010; Wang et al., 2011), galactose (Siebum et al., 2006; Turner et al., 2011), lactate (Gao et al., 2011), glycolate (Das et al., 2010), cholesterol (Pollegioni et al., 2009; Saxena et al., 2011), L-amino acid (Schrittwieser et al., 2011), D-aminoacid (Pollegioni & Molla, 2011) and monoamine oxidase (Buto et al., 1994; Edmondson et al., 2014). It is also plausible assuming that high ROS production in tadpoles in the CNF-I and CNF-II groups is associated with inflammasomes activation [intracellular multiprotein complexes activating caspases] by nanomaterials, whose reactive species formed in these systems are part of biochemical signaling reactions that can also activate inflammation through the production of several pro-inflammatory cytokines [see more details in Tschopp & Schroder (2010)]. Increased TBARS, mainly in the CNF-II group (Figure 4), suggested lipoperoxidation oxidative stress induced by CNFs, whose changes in biological membranes can further intensify ROS production (Itri et al., 2014).

Nevertheless, oxidative stress increase can cause different physiological consequences in organisms, such as increase in apoptotic and necrotic processes, as observed in our study (Figure 6). These data are particularly interesting, since they corroborate other studies that have already shown the induction of cell death processes in different model systems exposed to CNFs (either *in vitro* or *in vivo*). This finding is indicative of nanomaterials activating apoptotic and necrotic pathways through different pathways (Bottini et al., 2006; Elgrabli et al., 2008; Ravichandran et al., 2009; Patlolla et al., 2010; Srivastava et al., 2011; Wang et al., 2012; Kim et al., 2014; Salehcheh et al., 2020). Furthermore, data in our study also suggested that the increased rate of apoptotic and necrotic erythrocytes may have happened because of damages to cell membranes caused by direct contact of these models with CNFs or by increased oxidative stress, which was inferred through different biomarkers (H2O2, ROS and TBARS). However, assumingly, CNFs induced increased expression of apoptosis genes (as demonstrated by Lee et al (2015)], mitochondrial membrane potential collapse (as suggested by Salehcheh et al. (2020)], DNA damage [whose plausibility has already been demonstrated by Li et al. (2005) and Zhu et al. (2007)], and caspase activation [as suggested by Sohaebuddin et al. (2010)]. Shen et al. (2010) and Wang et al. (2012) have reported that CNs can cause, Ca2 + homeostasis imbalance and mitochondrial damage, as well as oxidative stress. These factors can be involved in MWCNTs-induced apoptosis and activate the production of the tumor necrosis factor by activating macrophages and monocytes, whose association with apoptosis and necrosis induction is well documented (Laster et al., 1988; Larrick & Wright, 1990; Van-Herreweghe et al., 2010; Ni et al., 2016; Yao & Cadwell, 2020; Liu &Jiao, 2020).

Interestingly, we also noticed neurotoxic effect on tadpoles exposed to CNFs, and this finding was mainly inferred through increased AChE and BChE activity (Figure 7). BChE played important role in supporting AChE in cholinergic transmission regulation, mainly in the absence of AChE (Li et al., 2000). However, these data are different from those reported in previous studies, such as those by Wang et al. (2007), Wang et al. (2009) and Cabral et al. (2013). Wang et al. (2009) and Cabral et al. (2013) reported that the anticholinesterase action of CNTs can be related to different action mechanisms, including the ones related to these nanomaterials’ ability to adsorb AChE, to compete with AChE for its substrate and even to reduce the higher reaction speed (V_max_) of this enzyme due to the substrate’s inability to reach the active site of the enzyme by immobilizing the nanomaterials. Wang et al. (2007) suggested that high BChE adsorption by the tested CNs promoted structural and functional changes that have led to significant reduction in enzyme activity.

Our data is following the study by Ibrahim et al. (2013), according to which, the direct effect of CNs on AChE activity did not cause significant change in the association and catalysis mechanism was observed. According to these authors, the catalytic constant increased as the Michaelis constant slightly decreased, and this finding is indicative of enzyme efficiency increase due to increased substrate affinity with the active site. The thermodynamic data of the enzyme’s activation mechanisms showed no change in substrate interaction mechanism with the anionic binding site. Therefore, assumingly, similar mechanism could explain the AChE and BChE increase identified in tadpoles exposed to CNFs. Therefore, it is possible that the activation of these enzymes took place due to the indirect effects of CNFs rather than to the aforementioned process. Studies carried out *in vitro* have already shown that H_2_O_2_ strongly increased the AChE activity (Schallreuter et al., 2004; Garcimartín et al., 2017), and it reinforces the hypothesis that the high production of this reactive oxygen species has also stimulated the cholinesterase activity. It is plausible supposing interactions between CNFs and acetylcholine receptors, and that such interactions led to increased AChE and BChE synthesis for the decomposition of higher levels of this neurotransmitter. The hypothesis that the stimulatory effect of CNFs on the activity of these enzymes has been associated with positive regulation of the AChE and BChE genes due to the inhibitory effect of nanomaterials, but it needs to be tested in future studies.

We also observed that the exposure to CNFs seems to have affected populations of neuromats living in some regions of tadpoles’ bodies, although in a different way. These cells are found in different amphibian species (Russel, 1976; Krupa et al., 2020) and make up a mechanosensory lateral line system with hair cells sensitive to movement, vibrations, and pressure gradients in the surrounding water (Lannoo, 1999). These cells are similar in morphology and function to hair cells in the auditory and vestibular system of other vertebrates (Mogdans, 2019; Roberts et al., 1988). Small movements in the water move the hair bundles of neuromast hair cells, and it mechanically opens the blocked ion channels (Harris et al., 1970; Sand et al., 1975). Hair cells (inside the neuromasts) depolarize and release neurotransmitters to the afferent neuronal terminals after water-flow deflection. These terminals transmit this information to the posterior brain (Jung et al., 2020).

Animals exposed to nanomaterials had fewer neuromast in their head (anterior) and a larger number of them in their caudal (posterior) region (Figure 8). This finding suggested differentiated action by CNFs, and it could have had important biological consequences in the evaluated animals. Neuromats in the head (be it in amphibians or in fish) are sensitive to surface wave movements in water, to detect prey, as well as present better spatial solution due to their greater density. Caudal neuromats (i.e., posterior) are more adept to detecting predators and water disorders (Russell, 1976; Schwartz & Hasler, 1996; Bleckmann & Zelick, 2009). Previous studies have also shown effect like that observed in our study, given differences between innervations of anterior and posterior neuromats. Hernandez et al. (2006) exposed *Danio rerio* larvae to different copper concentrations and reported differential hair cell regeneration between neuromats in the head and body of these larvae. Neuromats in the body were unable to regenerate at concentrations higher than 3.18 mg/L, whereas neuromats in the head regenerated at copper levels up to 25.42 mg/L. Similarly, posterior neuromasts were more sensitive in D. *rerio* embryos exposed to caffeine, dichlorvos, 4-nonylphenol and perfluorooctane sulfonic acid (Stengel et al. 2017). Posterior neuromats were more affected by copper sulphate and neomycin than previous neuromats in the aforementioned species after 30-min and 96-h exposure. Anterior neuromats exhibited greater cellular damage (Stengel et al. 2017). In this case, similarly to these findings, our data suggested differentiated action of CNFs on neuromats evaluated in the anterior and posterior regions of *P. cuvieri* tadpoles.

Although the action mechanisms of CNFs have not been explored in-depth in our study, it is tempting to speculate that these nanomaterials have acted in in neuromast populations through different ways. Assumingly, CNFs have affected these cells by competing with calcium ions at the fixation sites, and this process has avoided the flow of ions necessary for signal transduction, as observed by Hudspeth (1983) and Faucher et al. (2006). Thus, damage could be reversible. On the other hand, the reduced number of neuromats observed in the head of tadpoles exposed to CNFs can correspond to permanent damage to these cells because of increased oxidative stress, necrosis, or apoptosis. This hypothesis is supported by results of correlation analyses carried out between the number of neuromats in the head and CNF accumulation in the tested animals (see Table 1), as well as by reports by Olivari et al. (2008), who suggested similar mechanisms to explain the reduced number of neuromats in D. *rerio* larvae exposed to different copper concentrations. On the other hand, the increased number of neuromats observed in groups exposed to CNFs can be a physiological compensation mechanism to balance damages caused by nanomaterials to head cells, since the total number of neuromats did not differ between experimental groups (Figure 8C).

## 5. CONCLUSION

Based on the information above, our study confirmed the initial hypotheses and demonstrated that CNFs can accumulate in animals and have negative effects on the health of *P. cuvieri* tadpoles, even at short-term exposure, at environmentally relevant concentrations. The induction of nutritional deficit, oxidative stress and cyto-and neurotoxic effects are factors affecting these animals’ growth and development. However, it is necessary accepting that our results are only the “tip of the iceberg”; therefore, it is essential conducting further investigations to evaluate the biological impacts of CNFs on anurofauna. Limitations of our study are the starting point for future research. It is interesting further evaluating the long-term CBF’s effects and their impact on other physiological functions of the assessed model, as well as identifying and featuring possible damages caused by it in other amphibian species. This finding will be especially important to expand our knowledge about the action mechanisms of these pollutants. This information will be an important basis to assess ecotoxicological risks associated with the presence and dispersion of these pollutants in freshwater ecosystems and, their impact on anurofauna.

## 6. ACKNOWLEDGMENT

The authors are grateful to the Brazilian National Research Council (CNPq) (Brazilian research agency) (proc. N. 426531/2018-3) and to *Instituto Federal Goiano* for the financial support (Proc. N. 23219.000002.2021-81). The authors also acknowledge *Laboratório Multiusuário de Microscopia de Alta Resolução* (LabMic) (*Universidade Federal de Goiás/*Brazil) for their collaboration to the CNFs featuring process. Malafaia G. holds productivity scholarship granted by CNPq (proc, n. 307743/2018-7).

## 7. COMPLIANCE WITH ETHICAL STANDARDS

### Ethical approval

All experimental procedures were carried out in compliance with ethical guidelines on animal experimentation. Meticulous efforts were made to assure that animals suffered the least possible and to reduce external sources of stress, pain and discomfort. The current study did not exceed the number of animals necessary to produce trustworthy scientific data. This article does not refer to any study with human participants performed by any of the authors.

